# Estimating short-term synaptic plasticity from pre- and postsynaptic spiking

**DOI:** 10.1101/156687

**Authors:** Abed Ghanbari, Aleksey Malyshev, Maxim Volgushev, Ian H. Stevenson

**Affiliations:** University of Connecticut, Department of Biomedical Engineering; Institute of Higher Nervous Activity and Neurophysiology, Russian Academy of Science; University of Connecticut, Department of Psychological Sciences

**Keywords:** Short-term synaptic plasticity, Generalized Linear Model, Dynamical Synapse, Spike Statistics

## Abstract

Short-term synaptic plasticity (STP) critically affects the processing of information in neuronal circuits by reversibly changing the effective strength of connections between neurons on time scales from milliseconds to a few seconds. STP is traditionally studied using intracellular recordings of postsynaptic potentials or currents evoked by presynaptic spikes. However, STP also affects the statistics of postsynaptic spikes. Here we present two model-based approaches for estimating synaptic weights and short-term plasticity from pre- and postsynaptic spike observations alone. We extend a generalized linear model (GLM) that predicts postsynaptic spiking as a function of the observed pre- and postsynaptic spikes and allow the connection strength (coupling term in the GLM) to vary as a function of time based on the history of presynaptic spikes. Our first model assumes that STP follows a Tsodyks-Markram description of vesicle depletion and recovery. In a second model, we introduce a functional description of STP where we estimate the coupling term as a biophysically unrestrained function of the presynaptic inter-spike intervals. To validate the models, we test the accuracy of STP estimation using the spiking of pre- and postsynaptic neurons with known synaptic dynamics. We first test our models using the responses of layer 2/3 pyramidal neurons to simulated presynaptic input with different types of STP, and then use simulated spike trains to examine the effects of spike-frequency adaptation, stochastic vesicle release, spike sorting errors, and common input. We find that, using only spike observations, both model-based methods can accurately reconstruct the time-varying synaptic weights of presynaptic inputs for different types of STP. Our models also capture the differences in postsynaptic spike responses to presynaptic spikes following short vs long inter-spike intervals, similar to results reported for thalamocortical connections. These models may thus be useful tools for characterizing short-term plasticity from multi-electrode spike recordings in vivo.

**Author Summary:** Information processing in the nervous system critically depends on dynamic changes in the strength of connections between neurons. Short-term synaptic plasticity (STP), changes that occur on timescales from milliseconds to a few seconds, is thought to play a role in tasks such as speech recognition, motion detection, and working memory. Although intracellular recordings in slices of neural tissue have identified synaptic mechanisms of STP and have demonstrated its potential role in information processing, studying STP in intact animals, especially during behavior, is experimentally difficult. Unlike intracellular recordings, extracellular spiking of hundreds of neurons simultaneously can be recorded even in behaving animals. Here we developed two models that allow estimation of STP from extracellular spike recordings. We validate these models using results from in vitro experiments which simulate a realistic synaptic input from a population of presynaptic neurons with defined STP rules. Our results show that both new models can accurately recover the synaptic dynamics underlying spiking. These new methods will allow us to study STP using extracellular recordings, and therefore on a much larger scale than previously possible in behaving animals.

## Introduction

Short-term synaptic plasticity (STP) refers to fast and reversible changes of synaptic strength caused by the recent history of presynaptic spiking activity [1]. STP occurs on timescales from milliseconds to tens of seconds, and includes mechanisms for both facilitation of transmitter release, where synaptic strength increases with consecutive presynaptic spikes, and depression, where synaptic strength decreases. Facilitation and depression are mediated by the dynamics of presynaptic calcium and the depletion and replenishment of vesicles in the presynaptic terminals [1]. The relative contribution of facilitation and depression varies across synapses, cell types, and brain regions [2,3] with facilitation dominating at some synapses and depression at others. By shaping postsynaptic responses evoked by trains of presynaptic action potentials, STP alters neuronal information processing [4-6]. In vitro studies have shown that STP has profound effects on temporal filtering [7], network stability [7], and working memory [8]. Moreover, there is bidirectional interaction between STP and long-term synaptic changes: STP can determine the magnitude of long-term plasticity [9-12], and long-term synaptic changes also modify STP [12,13]. This results in an interplay between STP and long-term plasticity on multiple timescales [13,14]. Therefore, characterization of short-term plasticity in different systems is crucial for understanding neural computations.

Traditionally, short-term plasticity is studied using intracellular recordings where responses of the postsynaptic neuron to presynaptic stimulation are directly measured as evoked postsynaptic potentials or currents. Based on results of intracellular recordings Tsodyks, Markram, and colleagues developed a computational model that describes STP in terms of dynamics of resources and their utilization [15,16]. The Tsodyks-Markram (TM) model provides a phenomenological description of the short-term dynamics of synaptic responses in terms of 1) changes in the probability of transmitter release (utilization), related to the dynamics of presynaptic calcium and, 2) the use and replenishment of synaptic vesicles (resources). The TM model accurately captures the dynamics of synaptic responses caused by STP, links the observed diversity in synaptic dynamics to differences in the model parameters (utilization, recovery of resources, and their time constants), and allows prediction of postsynaptic responses to an arbitrary sequence of presynaptic stimuli [17]. Although several alternative models of STP have been proposed [18,19], the TM model is the most broadly used because it provides a compact description of STP with biophysically relevant parameters.

The TM model had been successfully used in a number of intracellular studies to assess synaptic dynamics in different connections [17,20,21] and changes of synaptic dynamics induced by long-term plasticity [13], adaptation [22] or injury [23]. Traditionally TM model parameters are estimated from responses to presynaptic stimuli applied in bursts of different frequencies [13,16,22,23]. A recent study presented a Bayesian approach that estimates TM model parameters by fitting postsynaptic responses induced by stochastic trains of presynaptic spikes [17]. Thus, STP parameters can be extracted from responses to in vivo-like presynaptic activity. Here we ask whether it is possible to estimate STP parameters using only the spike trains of pre- and postsynaptic neurons without access to postsynaptic potentials or currents. If available such a method would greatly expand the possibilities for studying STP in vivo. Although multiple intracellular recordings or simultaneous extra and intracellular recordings in vivo are possible [24-27], they are technically prohibitive for large-scale studies. Techniques for large-scale extracellular recordings, on the other hand, allow simultaneous recording of spiking from hundreds of neurons [28-30]. Prior studies compared cross-correlograms calculated using presynaptic spikes occurring after short or long inter-spike intervals, and found evidence for both short-term facilitation [31] and depression [32,33] of synaptic transmission in vivo. This split-correlograms approach, however, does not allow for a detailed reconstruction of synaptic weight for each presynaptic spike or estimation of underlying release probability and vesicular resources.

Here we develop two statistical methods that use pre- and postsynaptic spike trains to estimate the dynamics of short-term plasticity. Both approaches are based on a generalized linear model (GLM) that predicts postsynaptic spiking as a function of the observed pre- and postsynaptic spikes [34-38]. In these GLM-based methods we allow the effect of the presynaptic spikes to vary on short timescales as a function of the presynaptic spike timing. In a first model, the effect of presynaptic spikes is determined by the nonlinear dynamical equations of the TM model (TM-GLM). In a second model, we introduce a functional description of short-term plasticity based on a generalized bi-linear model (GBLM). Although the parameters in the second approach are no longer linked to biophysical properties the GBLM allows us to capture a wide range of neuronal interactions and synaptic dynamics.

To validate our models, we recorded spike responses of pyramidal neurons in vitro (cortical slices, layer 2/3 pyramids) to intracellularly injected currents composed of synaptic inputs with the known pre-defmed short-term plasticity. We show that, using only pre- and postsynaptic spike trains, the TM-GLM can recover the underlying parameters of STP, and the GBLM is able to reconstruct synaptic dynamics using a descriptive plasticity “rule”. Estimates provided by each of the two models were in good correspondence to ground truth values for a wide range of synaptic weights and time scales of facilitation and depression. Additionally, using simulated neurons we show that estimation of STP by these models is robust to several potential confounds: spike frequency adaptation, noise from probabilistic vesicle release, and spike sorting errors. The methods developed here, thus, have the potential to serve as powerful tools for large-scale studies of short-term synaptic plasticity in vivo, including alterations of short-term plasticity during different behaviors, during learning, or as a result of pathology.

## Results

Here we develop two model-based approaches to estimate short-term plasticity (STP) from trains of pre- and postsynaptic spikes. Both approaches are based on a generalized linear model (GLM) that predicts postsynaptic spiking as a function of the recent history of presynaptic spikes and the postsynaptic spikes. In the conventional GLM, the effect of presynaptic spikes is constant. In the new models we introduce a time-varying coupling term that depends on the history of presynaptic spikes and captures the short-term plasticity of synaptic connections.

In the first model, the coupling term is assumed to vary according to a Tsodyks-Markram model (Fig. 1, TM-GLM). The TM model provides a comprehensive description of STP using 4 physiologically motivated parameters: the baseline utilization of resources (U), the magnitude and time constant of facilitation (f and F), and the time constant for the recovery of resources (D). The dynamics of the synaptic resources and their utilization are described by two coupled differential equations that determine how postsynaptic responses depend on the history of presynaptic activity (Eq. 3 in the Methods). Using pre- and postsynaptic spike trains, the TM-GLM estimates both traditional GLM parameters (influence of postsynaptic spiking and coupling between pre- and postsynaptic activity) and the parameters ***θ = {D, F, U, f*}** describing short-term plasticity in the TM model.

**Fig 1:**
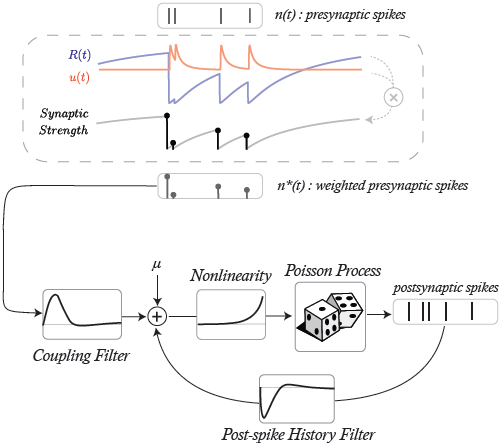
The output of the Tsodyks-Markram model provides input to a generalized linear model (GLM) where the weighted presynaptic spikes n*(t) combines with the history of the postsynaptic spiking to predict future postsynaptic spiking activity (bottom). The goal of our TM-GLM framework is then to estimate both the parameters of the TM (D, F, U, and f) and the parameters of the GLM (baseline firing rate, coupling filter, and post-spike filter) given only observations of pre- and postsynaptic spiking.

In the second model, we implement short-term plasticity as a descriptive rule which modifies the coupling term of the GLM based on specific presynaptic inter-spike intervals (ISIs). In this generalized bilinear model (GBLM, Fig. 2) the modification rule of the coupling term is not constrained by the known presynaptic mechanisms of short-term plasticity at unitary connections. However, the GBLM can still distinguish between facilitation (where presynaptic spikes following short ISIs have larger postsynaptic effects) and depression (where spikes following long ISIs have larger effects).

**Fig 2:**
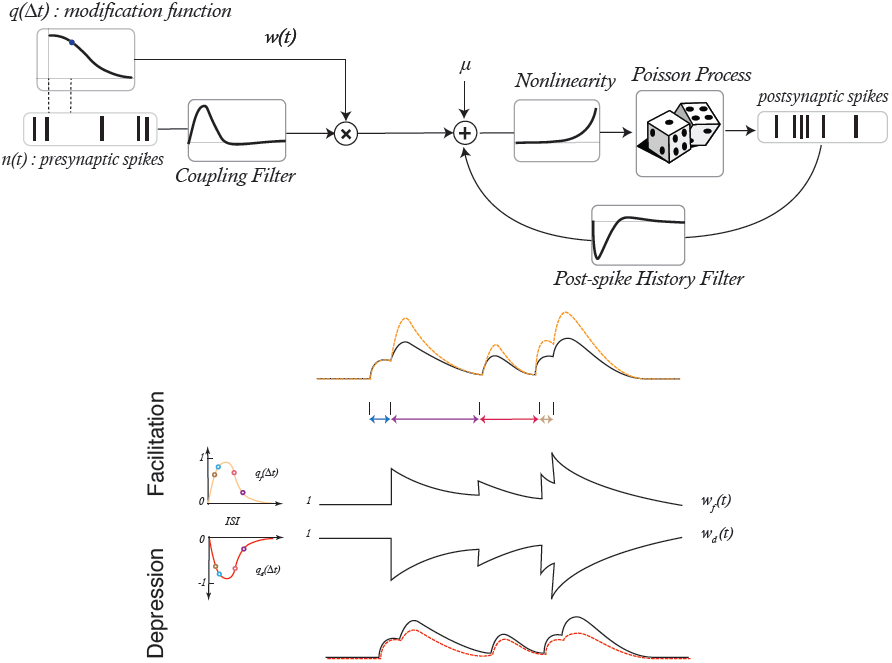
A generalized bilinear model (GBLM) provides a descriptive model of how synaptic weight varies as a function of presynaptic inter-spike intervals. The coupling term of a GLM is then weighted by a separate synaptic weight time-series w(t). w(t) is updated following presynaptic spikes according to a modification function q(Δt) and decays exponentially. When the modification function is positive the synaptic weight has facilitating dynamics, and when the modification function is negative the synaptic weight has depressing dynamics (bottom). The goal of our GBLM framework is to estimate the modification function (q) and the parameters of the GLM (coupling filter and post-spike filter) given only observations of pre- and postsynaptic spiking.

To validate the models, we examine how accurately they can reconstruct synaptic dynamics from spike trains of pairs of neurons connected by synapses with known plasticity rules. We obtained such data using the spiking of layer 2/3 pyramidal neurons evoked by injection of a fully-defined current generated by a population of simulated presynaptic inputs [39]. The advantage of these data is that they are generated by real neurons, with physiological spike generation mechanisms and post-synaptic dynamics. To examine the possible effects of additional factors that are present in in vivo recordings and may affect estimation of STP, we also used spike trains generated by simulated leaky integrate-and-fire neurons with: 1) spike frequency adaptation, 2) stochastic release at synaptic inputs, 3) spike sorting errors, and 4) correlated common input.

### Current Injection Experiments with Known Short-Term Synaptic Plasticity

To mimic recordings from pairs of neurons with known connectivity and short-term plasticity we made intracellular recordings from layer 2/3 pyramidal neurons in slices of rat visual cortex, and recorded spiking responses of neurons to injection of fully-defined fluctuating current. The injected current was designed to mimic the postsynaptic effect of synaptic inputs which have different strength and express unique synaptic dynamics (Fig. 3). To synthesize the current, we used a population of 96 presynaptic neurons where the spike times of each neuron were generated using an inhomogeneous Poisson process with a mean rate of 5 Hz. Six pools of 16 neurons (8 excitatory and 8 inhibitory in each pool) expressed five distinct types of STP, each defined by a unique set of parameters and ranging from strong depression to strong facilitation, along with a sixth pool of neurons which did not express STP (Table 1). STP of synaptic responses was implemented according to the TM model. Average synaptic weights for the 16 inputs in each pool ranged from strongly excitatory to strongly inhibitory, with excitatory and inhibitory inputs having the same amplitudes but opposite signs. This resulted in a balanced fluctuating current.

**Table 1:** The five parameter sets used to simulate presynaptic currents.

| TM parameters | D(s) | F(s) | U | f |
| --- | --- | --- | --- | --- |
| Strong Depression | 1.70 | 0.02 | 0.70 | 0.05 |
| Depression | 0.50 | 0.05 | 0.50 | 0.05 |
| Facilitation-Depression | 0.20 | 0.20 | 0.25 | 0.30 |
| Facilitation | 0.05 | 0.50 | 0.15 | 0.15 |
| Strong Facilitation | 0.02 | 1.00 | 0.10 | 0.11 |

**Fig 3:**
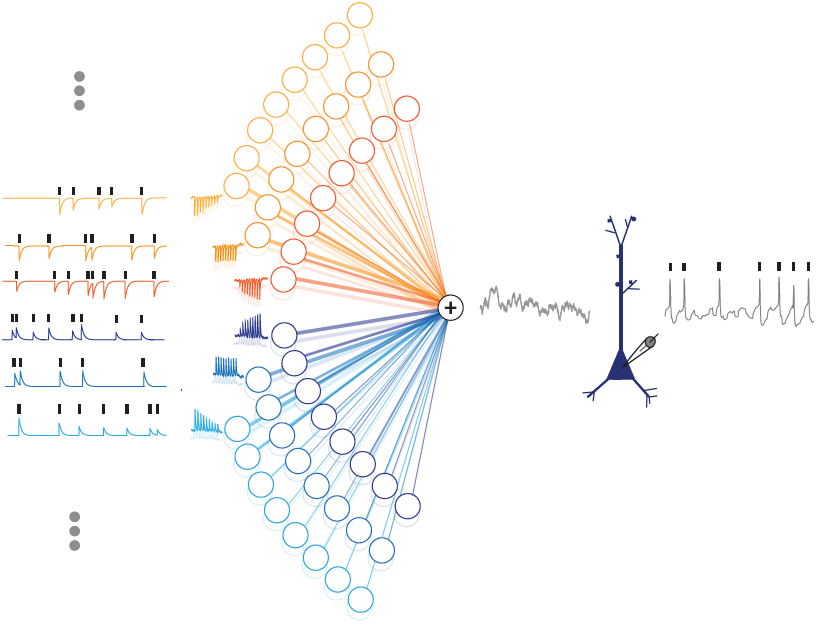
Artificial current injection to a Layer 2/3 pyramidal neuron. To validate our models, we recorded intracellularly from a cortical slice. We first simulated the spike times of 96 presynaptic neurons, then generated postsynaptic current traces corresponding to each input. Inputs had different types of plasticity ranging from strong depression to strong facilitation and a range of synaptic weights (both inhibitory and excitatory). The 96 current traces were then summed together and were intracellularly injected into the postsynaptic neuron whose spiking activity was recorded. These data then allow us to examine the relationship between pre- and postsynaptic spiking under 96 different plasticity/weight conditions.

Using the membrane potential responses to the injected current we detected postsynaptic spikes as positive-slope zero crossings. Thus, in this dataset we knew the timing of presynaptic spikes of each simulated presynaptic neuron, the time-varying synaptic weight, and the timing of the postsynaptic spikes.

To illustrate how STP at a single synapse affects postsynaptic firing in the presence of many other inputs, we performed a separate recording where the injected current had additional structure. One out of 96 presynaptic neurons repeatedly discharged with a pattern typically used for testing STP in slice experiments (9 regularly spaced spikes + 1 after a delay), while the spiking of the remaining 95 presynaptic neurons followed uncorrelated inhomogeneous Poisson processes as described above. This resulted in a repeating test pattern at one synapse embedded in fluctuating noise produced by the activity of the remaining presynaptic neurons. The strength of this synapse was increased to increase signal-to-noise ratio. The average postsynaptic current, membrane potential, and peristimulus time histogram of spiking (PSTH) in response to the test stimulation patterns demonstrate that the effect of a single strong input (>100pA) is clearly observable [Fig 4]. Moreover, synapses with different short-term synaptic dynamics: depression, facilitation and no plasticity produce distinct postsynaptic responses at all levels. In recordings with in vivo-like activity, the effects of short-term synaptic plasticity will be more subtle, since presynaptic spike times do not occur in such regular, repeating patterns under natural conditions and synaptic weights in neuronal connections are much weaker. The remaining analysis focuses on the recordings without the test patterns, where the strongest synaptic weights were -3 Op A.

**Fig 4:**
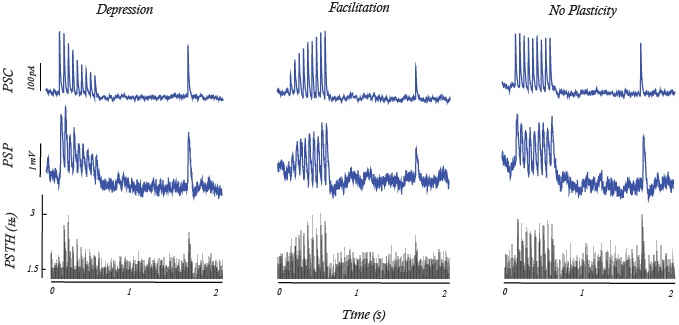
Postsynaptic responses to injection of a regular spike pattern immersed in fluctuating noise in vitro. To verify that the distinct types of STP directly affect postsynaptic spike statistics we compare the responses to a 50Hz train of presynaptic spikes for simulated synapses with short-term synaptic depression (left), facilitation (middle), and without plasticity (right). Only one out of 96 simulated presynaptic neurons had this regular activity pattern; the other 95 presynaptic neurons generated Poisson sequences of spikes to mimic the in vivo setting where postsynaptic neurons receive many presynaptic inputs (SNR=-17dB). Average of n=3000 repetitions shows clear effects of plasticity in the postsynaptic current and potential, but also in the postsynaptic spiking (PSTH).

In previous studies a split-correlogram approach had been used to reveal the effects of short-term plasticity on postsynaptic spike statistics in vivo [31,32]. By calculating cross-correlograms separately for presynaptic spikes following short ISIs (or in bursts) and for spikes following long ISIs (isolated spikes), evidence was found for both short-term facilitation [31] and depression [32,33] of synaptic transmission in vivo. To determine if this method of analysis could reveal effects of STP in our data obtained with inhomogeneous Poisson presynaptic spiking, we split presynaptic spike trains into spikes following inter-spike intervals shorter than the 10th percentile and longer than the 90th percentile of ISI distribution (Fig. 5A). Separate analysis of the postsynaptic effects of presynaptic spikes from these two groups revealed clear differences between synaptic inputs with distinct types of plasticity [Fig 5B]. In connections with depressing synapses the PSCs, PSPs, and, most importantly, peak spike counts in the correlograms were much reduced for short intervals. In connections with facilitating synapses, the postsynaptic effects were slightly increased following short presynaptic intervals [Fig 5B]. In synapses with intermediate forms of plasticity the effect of ISI on postsynaptic responses was less pronounced and was between the two extremes. Note that because of temporal summation after short ISIs the increase of the postsynaptic responses (PSCs, PSP, and spike count) is evident in the short interval correlograms even shortly before 0ms, similar to the results from in vivo study [31].

**Fig 5:**
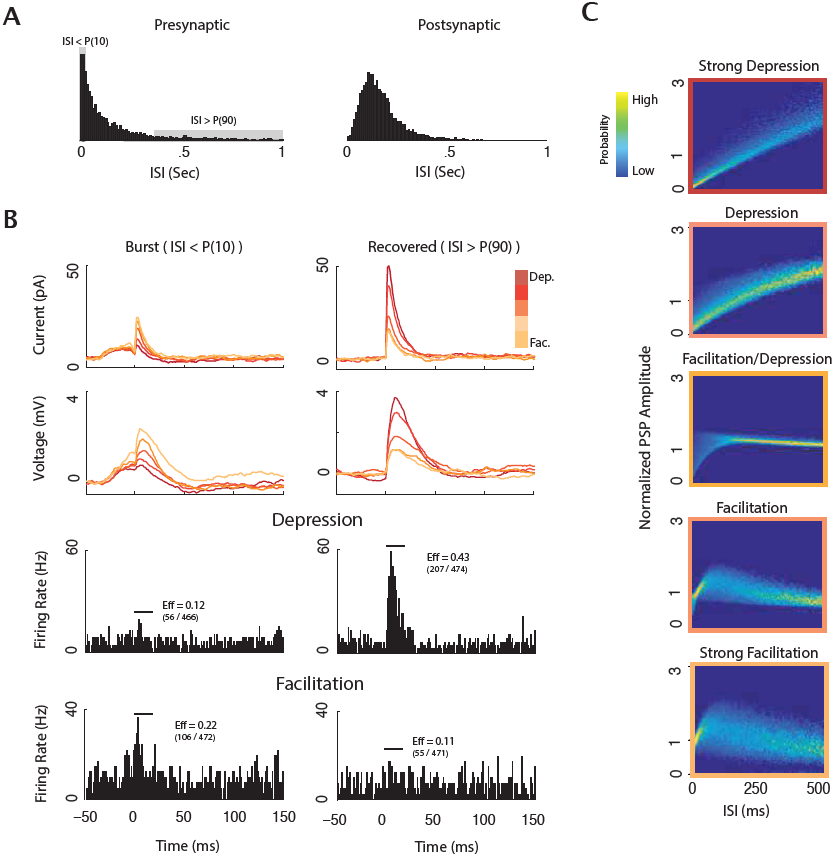
Postsynaptic responses for different presynaptic ISIs reveal the effects of STP. A) Inter-spike interval (ISI) distribution of a presynaptic and the postsynaptic neuron B) PSCs and PSPs [top] for strongest excitatory synapses for five different types of plasticity (colors) separated by presynaptic ISI (< 10^th^ percentile, left; >9 0^th^ percentile, right). Depressing synapses have much larger PSC/PSPs for ISIs >90^th^ percentile, while facilitating synapses have larger PSC/PSPs for the ISIs < 10^th^ percentile. Postsynaptic spiking [bottom] shows similar effects, but tend to be much more difficult to interpret due to the sparse spike responses. Split cross-correlograms are shown for two synapses: one with strong depression and one with strong facilitation [bottom]. For comparison, the PSPs and PSCs are vertically offset such that the average from -50ms to -30ms was set to 0. C) The distribution of PSP amplitudes as a function ofpresynaptic ISI for the different classes of plasticity used in experiment. One reason that split cross-correlograms are difficult to interpret is that there is no deterministic relationship between ISI and PSP amplitude, and, in some cases, such as with facilitating synapses, the relationship is nonlinear.

Thus, the effects of STP on spike responses of neurons to injection of a fully-defined current were clearly expressed in the difference between split-correlograms, consistent with results reported for in vivo recordings [31-33]. Our results show that the effects of ISI on split-correlograms were more pronounced for depressing than for facilitating inputs. One possible reason for such asymmetry may be that the presynaptic spike statistics used here does not fully elicit the effects of facilitation. To address this issue, we examined the distribution of PSP amplitudes as a function of inter-spike intervals for synapses with the different types of STP used in our model [Fig 5C]. While in depressing synapses the PSP amplitudes monotonically increase as ISIs increase, the response amplitudes in facilitating synapses depend on the ISIs in a nonmonotonic way. At facilitating synapses, there is an ISI range in which PSP amplitudes are elevated, but for both shorter and longer ISIs the amplitudes are reduced (Fig. 5C). This pattern makes it difficult to distinguish facilitation in split cross-correlograms, since short and long ISIs can produce similar PSP amplitudes. Moreover, facilitating responses also have higher variability than depressing responses for any given ISI, likely since stronger facilitation enhances the variability of utilization (release probability) compared with depressing synapses (Eq. 3). These factors appear to hinder detection of short-term facilitation with split-correlogram analyses.

The examples considered above show results for the strongest, excitatory simulated inputs (∼30 pA). Weaker excitatory synapses and inhibitory synapses express similar dynamics in their PSC and PSP amplitudes, however, the postsynaptic effects are less pronounced and show greater variability. For weak facilitating synapses there is often no detectable difference between the postsynaptic responses to short and long intervals. This analysis exposes a fundamental drawback of the split-correlogram approach: its low sensitivity to transient effects. By explicitly modeling how synapses vary in response to the history of presynaptic spiking, rather than modeling the average responses to only a single previous ISI, model-based approaches can more accurately reconstruct synaptic dynamics and distinguish between different types of STP.

### Inferring STP parameters from spike trains using the TM-GLM

We extend the GLM framework to include short-term synaptic plasticity implemented according to the Tsodyks-Markram model (see Methods). The TM model describes the dynamics of synaptic transmission using two coupled differential equations for resources *R* and their utilization (release probability) *u* with a set of four parameters ***θ = {D, F, U, f*}** (Eq. 3 in the Methods). To fit the TM-GLM to the observed spike trains we use an alternating coordinate ascent to maximize the (penalized) likelihood of observed postsynaptic spiking. Namely, we update the plasticity parameters with fixed GLM parameters and then update the GLM parameters with fixed plasticity parameters, alternating between the two optimization problems until the maximum is achieved. The TM formalism assigns a weight to each spike of the presynaptic neuron, while the GLM parameters characterize the influence of prior postsynaptic spiking and coupling between pre- and postsynaptic activity (as scaled by the TM weights). To facilitate convergence of the TM and the GLM parameters we impose prior constraints on both these parts of the model (see Methods). Using pre- and postsynaptic spike trains, we thus obtain estimates of both traditional GLM parameters and a complete set of parameters ***θ* = {*D, F, U, f*}** describing short-term plasticity in the TM model.

We fit the TM-GLM separately for each simulated connection in our in vitro recording. The 96 simulated presynaptic inputs had different weights and different types of STP, and our goal is to compare how these synaptic properties affect estimation of STP. Specifically, we have six sets of parameters corresponding to strong depression, depression, depression/facilitation, facilitation, strong facilitation, and a control set with no plasticity (Table 1). Although the optimization of the TM parameters is not convex, we find that, after adding informative priors (see Methods) the global optimum can be quickly found using random restarts. TM-GLM estimates of the time constant for depression *D* and the release probability *U* are closer to underlying true values than the estimates of the facilitation time constant *F* and its magnitude *f*. Fig. 6A shows results of bootstrapping to estimate the parameter uncertainty for the different types of plasticity. Note that high variability in the estimation of facilitation parameters is not a specific drawback of our model, but represents a more general problem. Indeed, previous work showed that estimates of facilitation parameters were non-precise even when direct measurements of postsynaptic responses, PSPs or PSCs (and not postsynaptic spikes as used in our model) were fitted [17,22]. Particularly for depressing synapses (where *U* is large and *F* is small), the estimation of *f*. is not well-posed. In this case, it may make more sense to use a more restricted TM model with fewer parameters [15,16] or to use a fully Bayesian approach where the posterior can be more completely assessed. More generally, the difficulty of estimating facilitation parameters might be a consequence of a relatively weaker effect of facilitation on postsynaptic activity as compared to depression. This interpretation is supported by the observation that despite the deviation of estimated parameters of facilitation from the true value, the model with the estimated parameters accurately predicts the steady-state filtering properties of dynamic synapses (Fig. 6C), as well as split cross-correlogram (Fig. 8A). Note that some of the bias in parameter estimation may be due to the choice of priors. Here we chose our priors to avoid local minima in the posterior that occur near the edges of parameter space, where *F* or *f*. are close to zero. However, as the number of observations increases these biases will be reduced, since likelihood will have a larger impact on the posterior than the prior. In general, the accuracy and confidence of the estimates will be affected by many factors, such as, the number and pattern of presynaptic spikes, number of postsynaptic spikes, the synaptic weight, and the type of STP.

**Fig 8:**
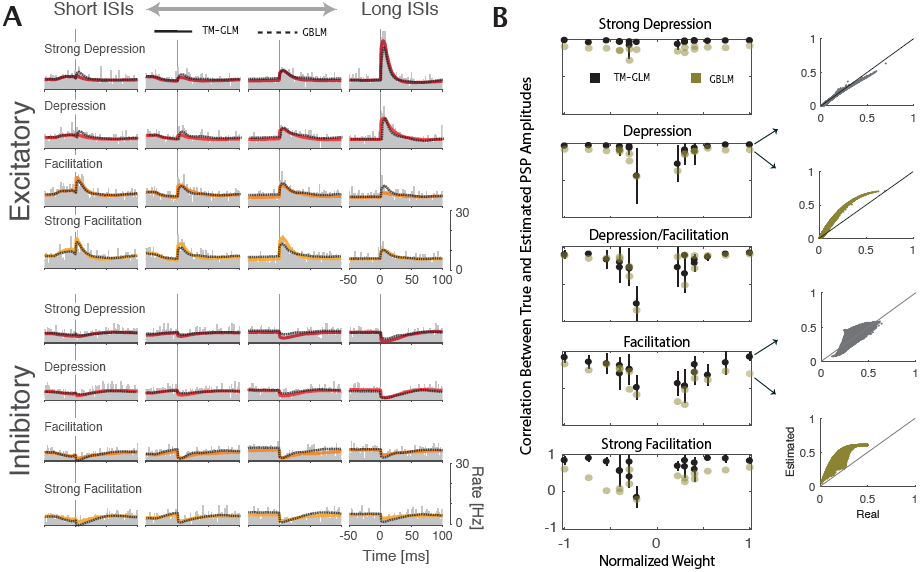
A) For each type of STP, the cross correlations between pre- and postsynaptic spiking split for the quartiles of the presynaptic ISI distribution (columns) from shorter (left) to longer inter-spike intervals (right). The estimated crosscorrelation from TM-GLM (solid) and GBLM (dashed) are shown on top of the observed cross-correlation (gray bars). B) The correlation between true and estimated amplitudes of postsynaptic potentials in five different classes of plasticity as a function of the overall synaptic (based on 1000s recording time with 5 Hz presynaptic firing rate). The PSPs of depressing synapses tend to be more accurately reconstructed than those of facilitating synapses, and weaker synapses (both excitatory and inhibitory) tend to be less accurately reconstructed than strong synapses.

**Fig 6:**
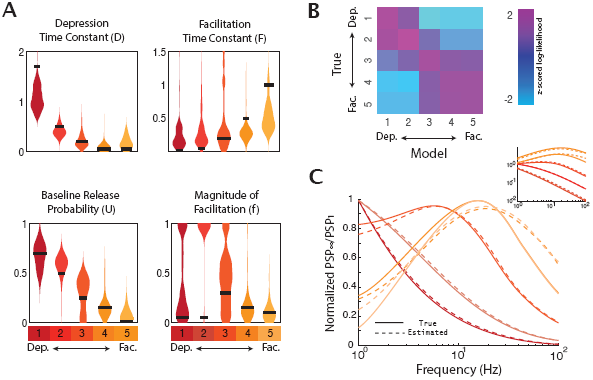
*Estimation of the TM parameters. A) Bootstrap distribution of the estimated parameters* **{***D,F, U,f***}** *for synapses with five different types of STP, from strong depression to strong facilitation (left to right). Here only the results for the strongest excitatory synapses are shown. The black horizontal bar in each distribution represents the true value for each artificial synapse (Table 1). B) z-scored log-likelihood values for strong synapses modeled with the parameters of each possible model. Even though the optimization of the TM-GLM is not guaranteed to be convex, we can accurately discriminate between different types of STP. Note that the highest likelihood is along the diagonal where the true type of STP corresponds to the same type modeled. C) Normalized steady-state postsynaptic potentials in response to a regular train of presynaptic spikes with different input frequency for true parameter sets (solid) and estimated parameters (dashed). Inset shows the unnormalized steady-state response on a log-scale.*

For large-scale analysis of STP in neuronal networks it might be important to distinguish between different types of plasticity at a synapse (e.g. facilitating vs depressing) and attribute certain types of plasticity to different classes of synaptic connections, rather than to extract the exact parameter values for each synapse. Again, although the problem is not convex, we find that the different types of plasticity can be distinguished based on spiking observations alone. For the 5 strongest excitatory inputs with each type of plasticity we compare the likelihood under the different settings of the TM parameters used in the recording [Fig 6B]. This analysis treats the problem of STP-identification as a classification problem. If the data do not provide a clear indication of the type of STP, e.g. for very weak synaptic inputs which have little effect on postsynaptic spiking, then the likelihood should be similar under all models - both facilitating and depressing. However, here we find that the true parameters do have the highest likelihoods, with depressing inputs having high likelihoods under the depressing model and facilitating inputs have high likelihoods under the facilitating model.

Additionally, even though the estimated parameters may differ from their true values, the (steady-state) synaptic dynamics of the estimated models typically matches the dynamics of the true models [Fig 6C]. Depressing synapses show characteristic low-pass filtering, while facilitating synapses have band-pass filtering with cutoff frequencies depending on the exact TM parameters.

### Inferring STP from spikes using a Generalized Bilinear Model

The TM-GLM estimates the short-term dynamics of a synapse described with biophysically realistic parameters that are related to the vesicle and calcium dynamics. In many cases, however, it might be useful to detach the description of the coupling between pre- and postsynaptic spiking from the biophysics of synaptic dynamics at an individual synapse. To describe neuronal interactions in terms of ISI-dependent modifications, we introduce a generalized bilinear model (GBLM, Fig. 2) that captures functional changes in the synaptic efficacy for different presynaptic intervals. In this model, the coupling term changes as function of presynaptic spiking, e.g. at facilitating synapses it increases for short ISIs, and at depressing synapses it decreases for short ISIs. We use basis splines to fit a smooth modification function (see Methods) that describes how the coupling term has been adjusted following different presynaptic intervals. We further assume that the effect of the modification is transient, decaying exponentially [Fig 2]. Compared to the TM-GLM, the GBLM has simplified description of the dynamics of coupling but provides a more explicit characterization of the effects of different ISIs on the modification of the coupling term.

The GBLM provides clearly distinct estimates of the modification functions for synaptic connections with different types of short-term plasticity [Fig 7]. For simulated inputs expressing the same type of STP, but having different weights (among strongest 3) or different signs (excitatory and inhibitory), the estimates of the modification functions were similar. These modification functions were estimated by maximizing the regularized log-likelihood. For stability, the spline basis was designed to have no effect on very short or very long ISIs where there is typically little data. However, for depressing synapses the modification function decreases the relative synaptic strength for ISIs between 0 and 1s, and for facilitating synapses the modification function increases the relative synaptic strengths.

**Fig 7:**
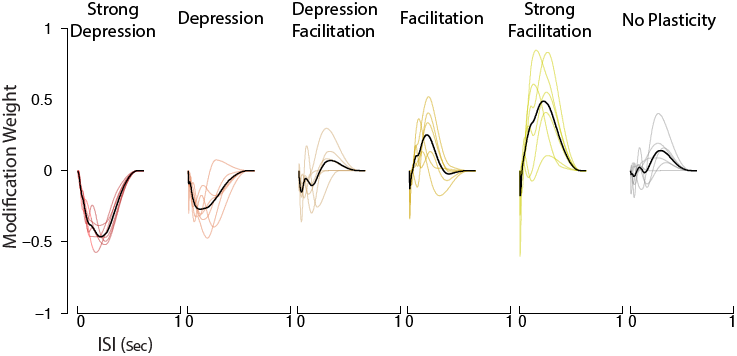
The modification function estimated using the generalized bilinear model (GBLM) for five different types of STP and for synapses with no plasticity. The modification functions for six strongest synapses (three inhibitory and three excitatory) are shown in color and the average is shown in black. These functions describe how synaptic weights change following different inter-spike intervals and allow the different types of STP to be distinguished. For strong depression, the modification function is negative and for strong facilitation the function is positive, capturing the respective decreases and increases in synaptic strengths.

### Comparison of the Models

Both the TM-GLM and the GBLM accurately describe split cross-correlograms for all examined types of STP, and for both excitatory and inhibitory inputs for the in vitro experiment [Fig 8A]. However, in addition to the spike statistics we can also compare how well the models reconstruct the time-varying individual PSC amplitudes. After estimating the plasticity dynamics for each simulated input using the TM-GLM 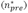 and the GBLM (*w*(*t*) ⊙ *n*_*pre*_) we then calculate correlations between the true PSC amplitudes and the estimated amplitudes under the two models [Fig 8B]. We find that the weights of the simulated inputs have a substantial effect on the reconstruction of PSC amplitudes. The estimated amplitudes at strong synapses (both excitatory and inhibitory) are reconstructed much more accurately than amplitudes at the weak synapses. Additionally, we find that the PSCs of depressing synapses are much more reliably reconstructed than PSCs of facilitating synapses (r=0.95±0.01 for synapses with strong depression vs. r=0.34±0.06 for synapses with strong facilitation). This is consistent with our observation that the PSCs of depressing synapses are more reliably related to ISIs compared to facilitating synapses [Fig 5]. Finally, the TM-GLM model appears to consistently out-perform the GBLM (average correlation for the TM-GLM across all types of plasticity and weights is r=0.70±0.03 compared to r=0.52±0.03 for the GBLM).

### Potential problems in raw spike data that may confound estimation of STP

In vitro recordings of responses to simulated presynaptic spikes have the advantage that the postsynaptic spikes are generated by the biophysics of a real neuron. However, estimation of STP from spike trains recorded in the intact brain in vivo may be compromised by several additional factors, not considered in this controlled experimental setting. Below we will analyze possible effects of four such factors on STP estimation: spike frequency adaptation, stochastic release of transmitter, uncertainty of spike sorting, and correlated common input. To examine how these sources of variability may affect the estimation of short-term synaptic plasticity from spikes we simulated postsynaptic spike trains using leaky integrate-and-fire model neurons receiving synaptic inputs with defined STP in the presence of noise. For simplicity, we focus on model synapses with strong depression, strong facilitation, and no plasticity (Table 1).

### Spike Frequency Adaptation

One factor that affects postsynaptic firing is spike frequency adaptation. In particular, an after-hyperpolarization (AHP) current mediating fast spike frequency adaptation can change the pattern of postsynaptic firing and may act to mask the influence of presynaptic STP on generation of postsynaptic spikes. To test if our models can differentiate the effects of AHP currents (I_AHP_), which alter the dynamics of the postsynaptic neuron, from the effects of short-term synaptic plasticity, we simulated two leaky integrate-and-fire (LIF) neurons with and without an I_AHP_ [40] (see Methods). In response to a long depolarizing pulse, the LIF neuron without an I_AHP_ fires at a stationary rate. The LIF neuron with the I_AHP,_ on the other hand, rapidly adapts - with a firing rate peaking immediately after the depolarization onset and gradually decreasing to a lower steady-state. After stimulus offset the firing rate of the adapting LIF decreases below the pre-stimulus level [Fig 9A]. These effects are not due to synaptic dynamics but reflect the dynamics of the postsynaptic neuron itself.

**Fig 9:**
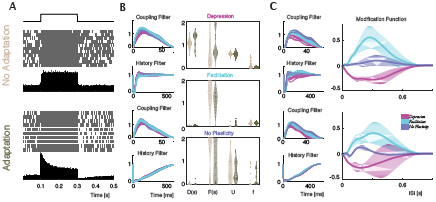
*Spike frequency adaptation affects post-spike history filters but does not affect STP estimation. A) Spiking of two LIF model neurons, with and without an* I_*AHP*_ *current, in response to a long depolarizing current step. Current step, spike rasters and PSTHs are shown for each model neuron. B) Parameters estimated by the TM-GLM. Estimated coupling filter and post-spike history filter of the two neurons in response to inhomogeneous Poisson inputs with short-time depression (blue), facilitation (red), and no plasticity (turquoise). Violin plots show estimated TM parameters for inputs with each type of the plasticity for the model neurons without (beige) and with* I_ahp_ *(green). C) GBLM estimates of coupling filter, post-spike history filter, and modification function for depressing (blue), facilitating (red), and no plasticity inputs (turquoise). Solid lines in the right panels show the average modification functions ±1 SD (bands).*

We simulated pre- and postsynaptic spike trains using the LIF model neurons (with and without an I_AHP)_ receiving inhomogeneous Poisson input with short-term synaptic dynamics governed by the TM model and applied our models to estimate STP from these spike trains. Results from the TM-GLM and GBLM for the two leaky IF neurons show that the adaptation properties mediated by I_AHP_ current are mostly captured in the post-spike history filters [Fig 9B-C]. For connections with depression, facilitation, or no STP, the estimated TM parameters and the modification functions estimated with the GBLM are similar with and without the I_AHP·_ Although frequency adaptation occurs on a similar timescale to shortterm synaptic plasticity, the methods here thus seem to be able to distinguish purely postsynaptic dynamics from the time-varying effect of the presynaptic neuron on the postsynaptic neuron.

### Stochastic Release

One further potential source of noise that is not included in the Tsodyks-Markram model, and that was not accounted for in our experiments in slices, is stochastic vesicle release. Although the TM model and the GBLM treat the synaptic transmission as deterministic and the PSC/PSP amplitudes can take any value, in real synapses PSC/PSP amplitudes are fundamentally stochastic with vesicles being probabilistically released from a limited number of sites. Compared to our in vitro experiments using the deterministic release, it may be more difficult to estimate STP parameters from the spiking of real neurons with stochastic release. To study how stochastic release impacts the estimation of STP parameters, we use a quantal model of synaptic plasticity [41,42]. In this model, the resources of the TM model are discretized based on the number of release sites and are then released according to a Binomial distribution with a time-varying probability given by the utilization variable of the TM model (see Methods). We simulated pre- and postsynaptic spike trains from LIF model neurons driven by inhomogeneous Poisson input with synaptic dynamics governed by the quantal TM model. The amplitudes of the postsynaptic currents are now noisy rather than deterministic functions of the presynaptic spike timing. In our simulations, increasing the number of release sites decreases the variance of the PSC amplitudes. For depressing synapses, stochastic release leads to a systematic bias in the estimates of the TM model parameters compared to their values under deterministic release [Fig 10A]. For facilitating synapses, on the other hand, the TM parameter estimation was not substantially affected. Similarly, the modification functions estimated with GBLM for depressing synapses were changed as the number of release sites is varied, while the modification functions for facilitation are more stable. Both the TM-GLM and GBLM can still distinguish between depression and facilitation, but considering stochastic release may be necessary for accurate parameter estimates in vivo.

**Fig 10:**
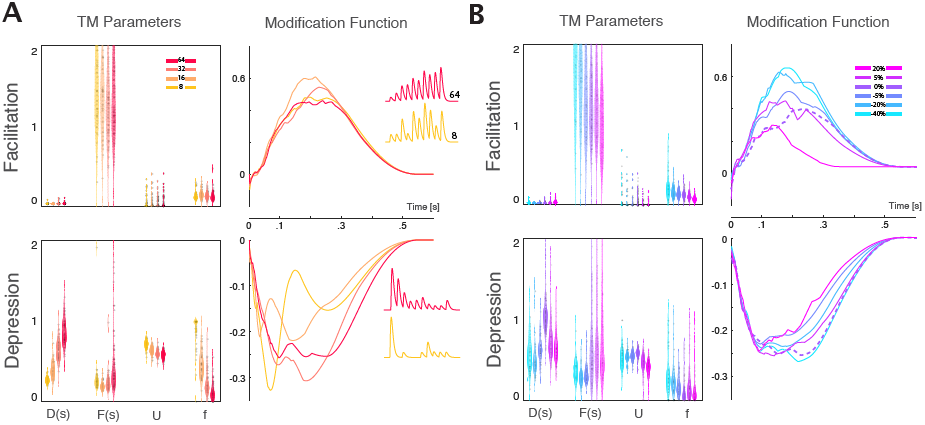
A) Stochastic vesicle release leads to a bias of STP estimates. Left panels; TM parameters for simulations with four different numbers of release sites: 8, 16, 32, and 64 (color coded). Right panels; median modification functions for 4 different release site numbers. Colors match with the left panel. B) Inaccurate spike sorting does not substantially alter STP estimates. Left panels; TM parameters estimated using simulated spiking of LIF neurons with random deletion or addition of spikes. From left to right: random deletion of -40%, -20%, -5% spikes; without deletion or insertion (0%), and with insertion of 5%, and 20%) of spikes. Right panels: median modification functions for the same sets with deleted and inserted spikes (colors match with the left panel). Dashed line denotes the estimated modification function under perfect spike sorting (without insertion or deletion).

### Spike Sorting

Another potential source of uncertainty, that may affect the estimation of synaptic dynamics from spikes, is imperfect spike sorting. In practice spike sorting from in vivo recordings is not a perfect process, and inaccuracies in spike sorting can lead to biased estimates of neural response properties [43]. Here, we simulated presynaptic and postsynaptic spike trains using LIF model neurons with strongly depressing or facilitating dynamics on inhomogeneous Poisson input (as above, See Table 1 for parameters). We then simulated the effects of imperfect spike sorting by randomly deleting and inserting spikes into both the pre- and postsynaptic spike trains before estimating STP. For insertion, we randomly selected spikes from two other inhomogeneous Poisson neurons (same baseline firing rates) and assigned the spikes to pre- and postsynaptic neurons. For both the TM-GLM and GBLM we find that the imperfect assignment of spikes (both addition and deletion) results in only small biases in the estimation of STP parameters for connections with strong facilitation and depression [Fig 10B]. Despite these small biases, we were able to distinguish between facilitation and depression even as the proportion of spike sorting errors becomes large (20-40% insertion/deletion).

### Common Input

In vivo, neurons often have common synaptic input from unobserved sources. Common input introduces correlations in pre- and postsynaptic spiking that are not due to synaptic connections between the recorded neurons. To study how such correlations would affect STP estimation we simulated a microcircuit with different levels of synchrony. In this simulation, two presynaptic neurons receive input from three sources: 1) a private, slowly fluctuating current, 2) a shared/common, slowly fluctuating current, and 3) an independent white noise current. The postsynaptic neuron receives the common input, an independent white noise current, and inputs from each presynaptic neuron - one with a depressing synapse and one with a facilitating synapse [Fig 11 A]. We then vary the strength of the common input using a weight parameter w, which determines how much of each neuron’s input is originating from the shared/common source and how much of the input comes from the private current. As the weight of the common input increases there is a short-term synchronization between the spiking of all neurons [Fig 11B]. At low *(w =* 0.25) and medium *(w =* 0.5) common input both the TM-GLM and GBLM were able to discriminate between depressing and facilitating inputs, but at *w =* 0.75 neither model was able to distinguish between the depressing and facilitating input. This simulation demonstrates that, at least in some situations, strong common input can cause both models to fail to estimate underlying short-term synaptic plasticity.

**Fig 11:**
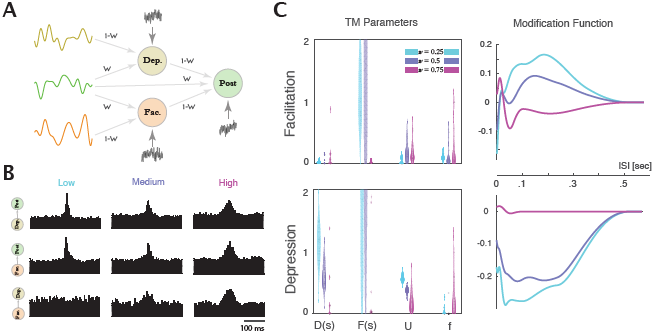
Common input can prevent accurate estimation of STP. A) We simulated a microcircuit where, rather than receiving independent input, the presynaptic and postsynaptic neuron both receive correlated, common input. We use three different sources of fluctuating input to each of the three LIF model neurons, varying the strength of common input. B) Cross-correlograms for three levels of common input (w= 0.25, 0.5, 0.75). C) Left panels: TM parameters estimated from spiking of neurons with (from left to right) low, medium, and high common input. Right panels: GBLM modification functions for the facilitating (top) and depressing (bottom) synapses and the three different levels of common input.

## Discussion

Intracellular recordings in brain slices have revealed a diversity of STP across cell types and anatomical connections [2,3]. Moreover, the details of STP at a given type of synapse may change depending on a multitude of factors, such as changes during development [44], neuromodulation [45], or induction of long-term plasticity [13]. Because STP critically affects information processing, understanding operation of neuronal networks during natural behavior requires large-scale analysis of STP in vivo. However, since large-scale intracellular recordings are not feasible in vivo, alternative methods are necessary for such studies. Large-scale extracellular recordings, on the other hand, are feasible in vivo. Existing techniques allow simultaneous recording of spiking of hundreds of neurons, and this number appears to be growing exponentially [28]. Characterizing short-term plasticity using spike observations is more difficult than using intracellular (PSC/PSP) signals, but short-term synaptic plasticity does have observable effects on spike statistics.

Prior evidence for STP in vivo obtained from spike trains alone employed a split cross-correlogram approach, in which the postsynaptic response to presynaptic spikes following short ISIs was compared to that following long ISIs. Several studies using this approach analyzed strong thalamocortical connections and found evidence for both short-term facilitation and depression [31-33]. To the best of our knowledge, however, the split cross-correlogram approach has not revealed evidence of short-term plasticity in weaker synapses, such as corticocortical connections. Here we introduce two new model-based methods to characterize short-term synaptic plasticity from pre- and postsynaptic spiking. By explicitly modeling synaptic dynamics these models are able to recover a detailed description of short-term plasticity. These models reproduce the results from split cross-correlograms (Fig 8), but also provide an explicit characterization of the dynamics of STP and allow reconstruction of PSP amplitudes for each presynaptic spike.

To validate our methods, we used spiking of layer 2/3 pyramidal neurons in vitro induced by injection of a current composed of PSCs from an artificial population of presynaptic neurons, whose spiking and plasticity parameters are known. Even though each presynaptic input represents only a small fraction of the total injected current, we can accurately estimate the synaptic dynamics from pre- and postsynaptic spiking. In this setting, both model-based methods, the TM-GLM and GBLM, can robustly distinguish between different types of STP, and can reconstruct PSP amplitudes for a wide range of synaptic weights for both excitatory and inhibitory connections. The TM-GLM provides a compact description of STP with four parameters related to the vesicular release and calcium dynamics in the presynaptic terminal. The GBLM provides a functional description of how the synaptic weight changes as a function of presynaptic ISIs. An advantage of the GBLM approach is that the synaptic modification rule is not constrained by the biophysics of single synapses, but has the potential to capture more complex dependences, including polysynaptic effects. One further advantage of the GBLM over the TM-GLM model is that the synaptic dynamics are assumed to be linear, which increases both the speed and robustness of the optimization process. Depending on whether a functional or a biophysical description is required, the two methods may thus both be useful tools for large-scale characterization of short-term synaptic plasticity from spiking activity.

Estimating synaptic plasticity from in vivo multi-electrode recordings of spiking activity will introduce several additional challenges. One challenge is simply detecting the connections between neurons. Strong monosynaptic connections are typically expressed in cross-correlograms as clear peaks (or troughs, for inhibition) with short latency and sharp onset, but weak connections or connections between neurons with low firing rates are difficult to detect in cross-correlograms. In previous work, we showed that model-based approaches can increase the sensitivity of detection for weak connections compared to traditional cross-correlation approaches [46], and the GLM-based approaches here are likely to have similar advantages.

A second challenge is that short-term synaptic plasticity isn’t the only source of variation in the observed postsynaptic responses to presynaptic spikes. Changes in the excitability of the postsynaptic neuron, stochasticity of vesicle release, and spike sorting errors can alter the statistics of the response and could potentially bias our estimates of short-term synaptic plasticity. To study how these sources of variability affect estimation of STP parameters we simulated spike trains of connected leaky integrate-and-fire model neurons, and introduced each of these confounding variables individually. We found that adding an after-hyperpolarization current (I_AHP)_ to the postsynaptic neuron impacts only the post spike-history filters in both the TM-GLM and GBLM, and does not substantially change STP estimation. Stochastic vesicle release and spike sorting errors, on the other hand, lead to biases in the estimation of short-term synaptic plasticity for our models. However, even with these additional noise sources, both the TM-GLM and GBLM are still able to reliably distinguish between connections expressing short-term facilitation and depression.

A third challenge is that correlations between the spiking of two neurons may be produced by common input rather than, or in addition to, the synaptic connection between the neurons. In our experiments with current injection in neurons in slices, inputs were generated as independent inhomogeneous Poisson processes, without the correlations that are present in vivo. To understand how correlated spiking can affect STP estimation, we simulated a small, feed-forward network of neurons with common input. We found that as the common input becomes stronger, the synchronization between pre- and postsynaptic spikes can interfere with the estimation of STP. The TM-GLM and GBLM were able to estimate synaptic dynamics only when common input was weak, but failed to accurately estimate the underlying synaptic dynamics for neurons with strong common input. While our in vitro experiment and simulations allowed us to compare STP estimation under controlled conditions with known synaptic dynamics, more work may thus be needed to account for all the dependencies that occur between pre- and postsynaptic neurons in vivo.

Finally, a fourth challenge is that the assumptions of the TM model itself do not necessarily describe the dynamics of all interactions between the pre- and postsynaptic neurons. The TM model only aims to describe presynaptic mechanisms of STP. However, postsynaptic factors such as desensitization or saturation of postsynaptic receptors may play a role in STP at some synapses, and the synaptic weight may vary on other timescales (e.g. due to LTP/LTD). Replacing the TM model with alternative models of plasticity may be a tractable approach to address these challenges [18,47,48]. Alternatively, since the GBLM is not constrained by single-synapse biophysics, it may, in some cases, provide a more flexible first-order description of short-term dynamics, including those that are not well described by the TM model.

Rather than describing anatomical connectivity, the two model-based methods introduced here describe the plasticity of functional interactions between neurons. Many of the techniques that have been used to improve models of functional connectivity without plasticity can be used to improve the TM-GLM and the GBLM presented here. For instance, it may useful to model multiple inputs simultaneously or to include latent common input in the model [49-51]. More structured regularization techniques may allow more accurate reconstruction with smaller sets of data [52,53]. To improve models of synaptic dynamics it may be useful to consider additional timescales [19], a higher-order expansion of the ISI dependencies, or other types of plasticity occurring on longer time scales, such as spike-timing dependent plasticity [54-56]. Applying these methods in vivo may then allow us to characterize short-term plasticity during natural behavior and in larger populations than previously possible.

## Methods and Models

### Ethics Statement

All animal use procedures conform to the principles outlined in the Guide for the Care and Use of Laboratory Animals (National Institutes of Health publication no. 86-23, revised 1985) and were approved by the Institutional Animal Care and Use Committee at the University of Connecticut.

### A phenomenological generative spiking model of short-term plasticity

Approaches using generalized linear models (GLMs) have proved to be effective tools for estimation neuronal connections from spike train data [57-59]. The standard GLM assumes that the spike train is a binary sequence of observations, *m*(*t*), generated from a Poisson process. For a single pair of neurons, we model the conditional intensity, *λ* (*t*), of this process as a linear combination of a baseline firing rate *μ*, a contribution from the presynaptic neuron ***rx***_***t***_ and weighted contribution from the postsynaptic spike-history ***sy*_*t*_** passed through an exponential nonlinearity (Fig. 1)

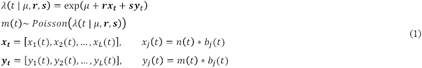

where *n*(*t*) and *m*(*t*) are the pre- and post-synaptic spike trains, respectively.

Our goal is to estimate the set of model parameters 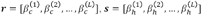 and *μ* describing the coupling, *k*(*t*), and post-spike filter, *h*(*t*), which best predicts the postsynaptic firing *m*(*t*).

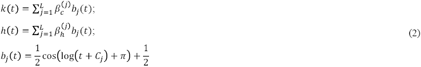

where *b*_*j*_(*t*) are raised-cosine basis functions which reduce dimensionality and allow a smooth representation of the two filters [35]. This stochastic model of a Poisson spiking neuron has a guaranteed convex log-likelihood which gives a unique set of parameters for its global maximum [38].

In order to model plasticity, we modified the GLM, allowing the contribution of coupling to vary over time. A conventional GLM treats all presynaptic spikes *n*(*t*), equally, with each presynaptic spike having the same “weight” when influencing conditional intensity, *λ* (*t*). To account for short-term facilitation and depression we modify the weights of each spike according to the phenomenological Tsodyks-Markram (TM) model [16]. The TM model describes the dynamics of resources R and their utilization U by the following system of differential equations:

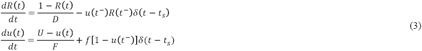

where resources, *R(t),* represent the portion of available vesicles which instantly decreases after each spike at *t*_*s*_ and gradually recovers with depression time constant *D.* The second equation describes release probability (utilization of resources), which instantly increases after each spike by *f*[1 — *u*(*t*^-^)], where *f* is the magnitude of facilitation and decays back to the baseline value, *U*, with facilitation time constant *F*.

The amplitude of the postsynaptic current *I*_*syn*_(*t*_*i*_) evoked by presynaptic spike at *U* is described by

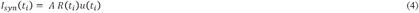

where *A* is the maximal current that can be evoked at that synapse if all resources are recovered *(R =* 1) and are released at

With different sets of parameters ***θ*= {*D,F,U,f*}** this model can reproduce diverse types of short-term plasticity (depression, facilitation or a mixture of both) observed experimentally [17]. Using these dynamics, we create a “marked” point-process

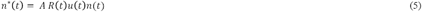

where *n*\*(*t*) captures the amplitudes of PSCs at the time-points of presynaptic spikes and is zero otherwise. By using *n*\*(*t*) instead of *n*(*t*) in the modified GLM (TM-GLM), we account for STP in the coupling term. Note that when *n*\*(*t*) is constant *(R(t)* and *u*(*t*) constant) the TM-GLM will describe a steady-state synapse with no short-term plasticity, and, in this case, TM-GLM is identical to the original GLM.

With the modified coupling term the original observation model is rewritten as

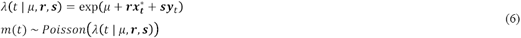

Using the TM-GLM our goal is to estimate the static parameters of the synapse *ϕ* = {*μ,****r,s***}, as well as the plasticity parameters ***θ*** = {***D, F, U,*** *f*}, given the pre- and postsynaptic spike trains. Specifically, we aim to find maximum *a posteriori* (MAP) estimates of *θ* and *ϕp* that optimize

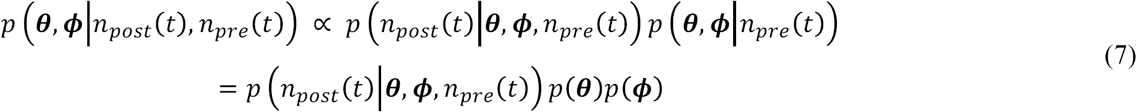

To prevent over-fitting and assure nonnegative values, we introduce weakly informative priors on the plasticity parameters *p*(*θ*) to span the parameters space only over meaningful intervals and prevent the optimization from getting stuck at local minima. We then use coordinate ascent, maximizing the log posterior by alternating between optimizing the plasticity parameters given the GLM parameters and fitting the GLM given fixed plasticity parameters. Although this posterior is not guaranteed to be convex, in many cases, the non-convexity of GLM-like models does not lead to optimization problems [60-62]. Previous work estimating STP parameters from intracellular recordings suggests that, rather than point estimates, a fully Bayesian approach may provide a more accurate understanding of the parameters [17,41]. Although it is possible to use MCMC to sample from the posterior, the large number of function evaluations (compared to optimization) makes it less attractive for our model with spike observations.

When optimizing plasticity parameters (GLM parameters fixed) we randomly restart over the 0-space and use priors {0 < ***D, F <*** 2} **∼** *gamma(α* ***=*** 1.2, *β* **=** 2) and {0 **< *U,f* <** 1} **∼** *beta(* 1.01,1.01). We optimize the plasticity parameters in the log-domain using two-metric projection and numerical differentiation of the posterior [63]. Additionally, we found that when optimizing the plasticity parameters, convergence is improved by normalizing the static coupling term *k(t*) and optimizing an amplitude *A* (with prior *p(A) = cauchy(*0,50)) alongside the parameters 0. These prior distributions and parameters were chosen to prevent the model from reaching the boundaries (e.g. *U* or *f* at 0 or 1), but they do introduce bias into the parameter estimates and may not necessarily work well for all sets of data.

When optimizing the static GLM parameters *ϕ* (plasticity parameters fixed) we would typically assume *p*(*ϕ*) to be flat. However, we found that in some cases the coupling term *k(t*) interacts with the plasticity parameters. For instance, an excitatory depressing synapse will show a biphasic coupling term where a negative component can partially account for the reduced impact of a burst. To prevent this type of ambiguity we introduced a quadratic penalty on negative coupling coefficients r with the improper prior log *p*(*r*) ∝ — *η r*^*2*^ **l**_*r*<0_. In practice, we use LBFGS optimization of the penalized log-likelihood and this ensures that the estimated coupling term is approximately positive for excitatory inputs and negative for inhibitory inputs. With limited data or when extending the model to multiple inputs additional types of regularization may be useful [58].

### A nonparametric generalized bilinear model of STP

The phenomenological model, described above, gives a clear view of the synaptic dynamics by searching over the θ -space of STP parameters. However, in cases were TM assumptions on synaptic dynamics such as vesicle release and changes of the calcium changes in presynaptic terminal doesn’t hold, it may be preferable to have a model of STP that is not constrained to the TM dynamics. In a second type of model - the generalized bilinear model - instead of searching over the space of STP parameters we directly infer a short-term synaptic modification “rule”. This generalized bilinear model (GBLM) compartmentalizes the coupling term into a stationary and a short-term plastic modification (Fig. 2).

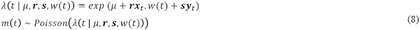

Here the modification term, *w*(*t*), weights the static coupling term depending on the history of presynaptic spiking. For a synapse with no plasticity, *w*(*t*) equals to one and the coupling term, ***rx***_*t*_ is static and does not depend on previous presynaptic spiking. For a synapse with plasticity, *w*(*t*) >1 would increase the static coupling term ***rx***_*t*_ to account for facilitation and *w*(*t*) <1 would decrease the ***rx***_*t*_ to account for depression. In both cases, the effect of w(t) on the coupling term decays with time and the coupling term recovers to its static form. We defined the modification function as:

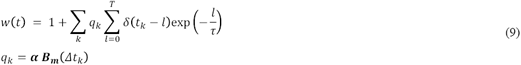

where q(·) determines the amplitude of exponentially decaying effects from previous spikes on the synaptic weight. Here *k* indexes the presynaptic spikes with times *t*_*k*_ and previous inter-spike intervals Δ *t*_*k*_.

Although we could attempt to fit the decay function (instead of using single exponential) and its time-constant (*τ* = .2) we fixed them to increase the robustness and speed of the maximum likelihood parameter fitting. Spikes are then convolved with the exponential kernel weighted by the modification terms *q* ((·). To ensure q((·) is a smooth function we represent it using the B-spline bases, *B*_*m*_*(Δt*), with log-spaced sampling knots in *Δt*_*k*_. The final model is linear in both the stationary parameters, *{μ,* ***r,* s**}, and STP parameters, ***q***. To estimate the parameters, we alternate between two GLMs: fitting {*μ,****r,s***} with fixed ***q*** and fitting ***q*** with fixed {*μ,* ***r, s***} (both using iterative reweighted least squares - IRLS). Although the two GLMs are log-concave in this problem, the joint likelihood of {*μ,* ***r, s, q***} is not guaranteed to be concave. However, we find that in practice convergence is fast using the alternating method and random restarts results in the same final solution.

### Experiments in Slices: Recording and Current Injection

Slices of visual cortex were prepared from male Wistar rats (P21-P23) as described in detail in our prior work [39]. Extracellular solution used during preparation of slices and for perfusion of recording chamber contained (in mM): 125 NaCl, 2.5 KC1, 2 CaCl_2_, 1 MgCl_2_, 1.25 NaH_2_P0_4_, 25 NaHC0_3_, 25 D-glucose and was bubbled with 95% 0_2_ and 5% C0_2_. Patch clamp electrodes for whole cell recordings were filled with K-gluconate based solution (in mM: 130 K-Gluconate, 20 KC1, 4 Mg-ATP, 0.3 Na_2_-GTP, 10 Na-Phosphocreatine, 10 HEPES) and had a resistance of 4-6 MH. Whole-cell recordings were made from layer 2/3 pyramidal neurons of rat visual cortex. Membrane potential responses to injection of fully-defined fluctuating current ([39]; see below) were recorded using the bridge mode of a Dagan B VC-700A amplifier (Dagan Corporation, USA). Data were digitized at 20 kHz (Digidata 1440A, Molecular Devices, USA) and stored for further processing. Timings of postsynaptic spikes were determined as positive-slope zero crossings of the membrane potential signal.

An artificial current for injection was designed to mimic the postsynaptic effect of a realistic cortical circuitry with inputs of different strength and unique short-term synaptic characteristics (Fig. 3). Current for injection was synthesized using a population of 96 presynaptic neurons (6 pools of 16 neurons, 8 excitatory and 8 inhibitory). Five sets of STP parameters were chosen to cover the whole spectrum of the plasticity from strong depression to strong facilitation. The sixth set of synapses did not express short-term plasticity. For each neuron, we generated an inhomogeneous Poisson spiking series with the log rate generated using a cubic spline function with 1 knot/s and standard normally distributed amplitudes. The rate is then scaled to generate an average spike rate of 5Hz and the spikes are weighted to generate the postsynaptic current amplitudes of the TM model. The weighted series of postsynaptic current amplitudes was then convolved with a synaptic integration kernel to generate the artificial postsynaptic current traces. We generated the kernel as a difference of two exponentials with time constants of 1ms and 10ms. Eight different synaptic weights with a normal inverse cumulative distribution function *{μ*= .7 & *σ =* .93) were used to create a pool of excitatory synapses. Same synaptic weights, but witha negative sign were used to generate currents produced by inhibitory neurons. Because the number and weight distributions for excitatory and inhibitory presynaptic neurons were the same, the total input current was balanced. We used 20 different realizations of the current for injection. The duration of each current trace was 46s. Injections of fluctuating currents were separated by intervals of 60-100s. The amplitude of the injected current was adjusted to produce membrane potential fluctuations of 10-15 mV. DC current was added to achieve the average postsynaptic firing rate of ∼5Hz.

Thus, we knew the timing of presynaptic spikes for each simulated presynaptic neuron contributing a synaptic connection as well as its amplitude and the parameters governing its short-term plasticity. We used individual pairs of pre- and postsynaptic spike trains to compare the parameters of short-term plasticity, estimated by the models, to the ground truth values.

### Simulation: Leaky integrate?and?fire model with adaptation

To examine the limitations of our models more thoroughly, we simulated a leaky integrate-and-fire model neuron receiving presynaptic input with short-term synaptic plasticity. In particular, to examine the effect of spike frequency adaptation we simulate a postsynaptic neuron with and without an after-hyperpolarization current [40]:

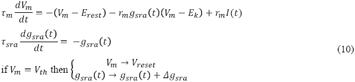

where *E*_*k*_ is the reversal potential due to K^+^, *g*_*sra*_*(t)* the spike rate adaptation conductance, which changes with rate *Ag*_*sra*_ — 200*nS*, and *E*_*k*_ *=* 80 *mV* is the reversal potential. The other parameters were set to *E*_*rest*_ *=* 80*m*7, *E*_*k*_ *=* 80*m*7, *V*_*rese*__*t*_ 80 ?*mV, V*_*th*_ *=* 54*m*7, *τ*_*m*_ = 10*ms, τ*_*sra*_ *=* 100*ms,* and *r*_*m*_ *=* 10*MΩ.* Similar to the in vitro experiment above, *I*(*t*) is synthesized by simulating a presynaptic input with short-term synaptic plasticity (inhomogeneous Poisson spiking with Tsodyks-Markram PSC amplitudes). We then adjust the DC current, noise, and synaptic strength to achieve the desired postsynaptic spike rate (5Hz) along with a cross-correlogram similar to those obtained by the strongest synapses in the in vitro experiment.

### Simulation: Stochastic model of short-term synaptic plasticity

Although the TM model treats short-term synaptic plasticity as a deterministic process, synaptic transmission is a discrete, stochastic process where a discrete number of vesicles are present and probabilistically released following presynaptic spikes. To model this additional variability, we use LIF simulations, as above, where rather than having PSC amplitudes be synthesized from the TM model we use a quantal, stochastic extension of the TM model.

First, to make the TM model discrete, we consider an integer number of release sites, *n*_*max*_, where, at any point in time, only a fraction of resources are available to be released, *a*_*m*_ *=* ⌊*n*_*max*_ *R*_*m*_ ⌋. Following each presynaptic spike, a discrete number of vesicles is released

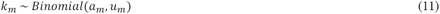

giving the PSC amplitudes

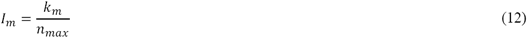

Following a spike, the resources and utilization at the following spike (after interval *Δt)* are given by

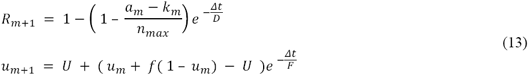

## Acknowledgements

We are grateful to Vladimir Ilin for help in setting up the experiments.

